# Cerebellar granule cell precursors can extend processes, undergo short migratory movements and express postmitotic markers before mitosis in the chick EGL

**DOI:** 10.1101/220715

**Authors:** Michalina Hanzel, Richard JT Wingate

**Affiliations:** MRC Centre for Neurodevelopmental Disorders, King’s College London, UK

## Abstract

Cerebellar granule cell precursors (GCPs) form a secondary germinative epithelium, the external germinal layer (EGL) where they proliferate extensively to produce the most numerous cell type in the brain. The morphological sequence of events that characterizes the differentiation of GCPs in the EGL is well established. However, morphologies of individual GCP and their differentiation status have never been correlated. Here, we examine the morphological features and transitions of GCPs in the chicken cerebellum by labelling a subset of GCPs with a stable genomic expression of a GFP transgene and following their development within the EGL in fixed tissue and using time-lapse imaging. We use immunohistochemistry to observe cellular morphologies of mitotic and differentiating GCPs to better understand their differentiation dynamics. Results reveal that mitotic activities of GCPs are more complex and dynamic than currently appreciated. While most GCPs divide in the outer and middle EGL, some are capable of division in the inner EGL. Some GCPs remain mitotically active during process extension and tangential migration and retract their processes prior to each cell division. The mitotically active precursors can also express differentiation markers such as TAG1 and NeuroD1. Further, we explore the result of misexpression of NeuroD1 on granule cell development. When misexpressed in GCPs, NeuroD1 leads to premature differentiation, defects in migration and reduced cerebellar size and foliation. Overall, we provide the first characterisation of individual morphologies of mitotically active cerebellar GCPs *in ovo* and reaffirm the role of NeuroD1 as a differentiation factor in the development of cerebellar granule cells.

## Introduction

Transit amplification of basal progenitors is an important feature of the vertebrate brain development that allows for a rapid expansion of specific cell populations and has facilitated the extraordinary evolution of foliated structures such as the cortex and the cerebellum^1–3^. Secondary proliferation also allows dedicated progenitors to respond to local environmental conditions to populate neural structures as required during development and repair. The process is found, for example, in the progenitors in the subventricular zone (SVZ) that generate the migrating neuroblasts of the rostral migratory stream (RMS)^4^, the basal neocortical progenitors^1,5,6^, and the progenitors in the external germinal layer (EGL) of the cerebellum. In addition to potentiating exponential growth, the adaptive behaviour of secondary proliferative populations may also underlie developmental plasticity.

The most well studied secondary proliferative population in the brain is the EGL. The traditional view of EGL assembly, established by Cajal^7^, is that granule cell precursors (GCPs) born at the rhombic lip populate the dorsal cerebellar anlagen where they undergo determined sequential phases of proliferation, morphological elaboration, followed by tangential and radial migration into the inner granular layer (IGL) (Fig. 1A). This view has had a profound influence on how morphology and differentiation status of GCPs is interpreted and has been observed in many species and systems^8–18^, including chick cerebella^11^ (Fig. 1B). However, does this deterministic and linear interpretation of morphology capture the diversity of GCP behaviour? Given that the neuroblasts of the RMS, for example, retain their ability to divide as they migrate towards the olfactory bulb^19–22^ and express postmitotic markers^23–25^, we decided to explore the possible presence of similar developmental features in GCPs in the cerebellum.

**Figure 1.**
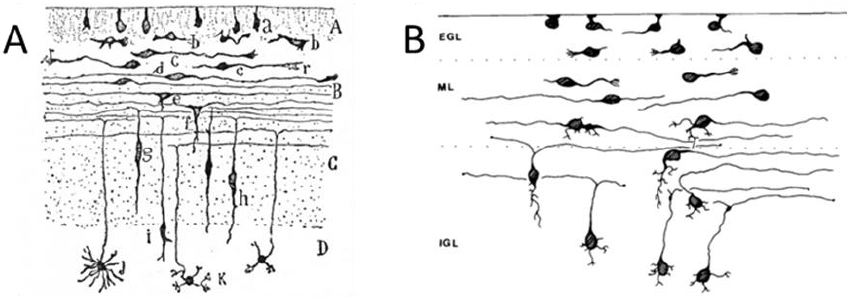
Schematics of cerebellar granule cell morphological transition in rats and chicken. **A**) Picture of granule cell morphological development drawn by Cajal based on Golgi staining of rat tissue. A, Layer of undifferentiated cells; B, layer of cells in horizontal bipolar stage; C, partly formed molecular (plexiform) layer; D, granular layer; b, beginning differentiation of granule cells; c, cells in monopolar stage; d, cells in bipolar stage; e,f, beginning of descending dendrite and of unipolarisation of cell; g,h, i, different stages of unipolarisation or formation of single process connecting with the original two processes; j, cell showing differentiating and completed dendrites; k, fully formed granule cell (caption from Bailey and Miller, 1921). **B**) Drawing of chick granule cell morphological differentiation based on a Golgi staining study (Quesada and Genis-Galvez, 1983). Morphological transitions of chick granule cells resemble closely those found in mammals such as the mouse and rat.

Given the number of studies of EGL development, there is a surprising dearth of time-lapse movies of GCP behaviour in an intact EGL, and the few performed do not explore progenitor morphology^26,27^. One reason for this could be the high density of cell packing in the EGL that makes visualisation of detailed morphology difficult. To overcome this, we electroporated a GFP transgene into chick rhombic lip cells in early development and allowed the cerebellum to grow *in ovo* for ten days. This resulted in sparse labelling of EGL cells and allowed detailed morphological examination of GCPs. We find, in fixed tissue and *ex ovo* time-lapse imaging of cerebellar organotypic slices, that GCPs retain their ability to divide in all layers of the EGL, are highly motile between cell divisions, elaborate long and complex cellular processes that are retracted prior to cytokinesis, and can express postmitotic markers before undergoing mitosis.

## Results

To observe the morphologies of individual GCPs in the chick EGL, we electroporated embryonic day 5 (E5) embryos with a plasmid encoding a GFP transgene flanked by Tol2 sites, and a plasmid encoding a Tol2 transposase, resulting in a stable genomic expression of GFP in a subset of granule cells born at the rhombic lip. At E14, the peak period of GCPs proliferation in the chick EGL, we sacrificed the embryos and examined fixed tissue with immunohistochemistry, as well as performed time-lapse imaging of the living cells in the EGL in organotypic cerebellar slices. We find that there are sparsely labelled rhombic-lip derived cells in the EGL at this stage, representing granule cells at various stages of development (Fig. 2A-B).

**Figure 2.**
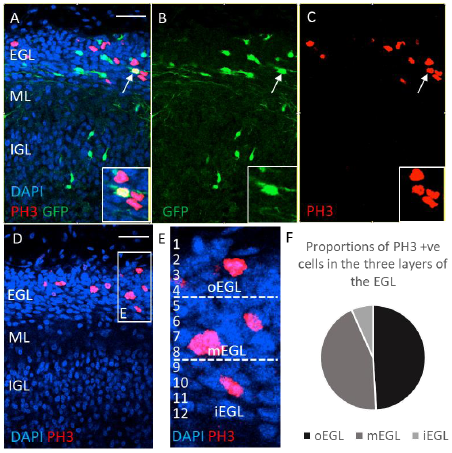
GFP-electroporated cells labelled with PH3 antibody are found in all layers of the EGL. Embryos were electroporated with Tol2:GFP at E4 and sacrificed at E14. Cerebella were sectioned in a coronal orientation and stained with anti-PH3 antibody. **A-C)** show an example of the cerebellar tissue where (**B**) GFP electroporated cells can be seen in the EGL. (**C**) PH3 expressing cells are shown in red. Some electroporated cells co-express PH3 (insert). Morphologies of cells in the different layers of the EGL are seen. In the inner layers cells have differentiated morphologies and long processes yet PH3^+ve^ cells are present among them. **D)** EGL of E14 embryos was divided into three equal layers and the numbers of PH3^+ve^ cells in each layer was quantified. PH3 is distributed in all layers of the EGL **E)** An example of how the EGL was divided into three layers. **F)** Numbers of PH3^+ve^ cells were quantified in the three layers of the EGL in 15 slices from 3 cerebella. Most cells were present in the outer EGL (49%), but nearly equally many were present in the middle EGL (44%). Considerably fewer cells were present in the inner EGL (7%). Scale bar= 100μm

To examine the precursors, we used antibodies against phosphohistone H3 (PH3) as a marker of proliferating cells in the EGL. There are no known cells other than GCPs that proliferate within the EGL at this stage. The EGL was divided into three equal layers (outer, middle, inner). As shown in Fig. 2B, PH3 positive cells were found in all layers of the EGL, including the inner EGL. The highest proportion of PH3 positive cells was found in the outer EGL (49%), followed by the middle EGL (44%) with a minority of cells located within the inner EGL (7%).

Proliferating GCPs are traditionally only expected to localise to the upper half of the EGL and ‘have a round soma without any long processes’^16^. We therefore characterised the morphologies of electroporated cells staining for PH3 located in the different EGL layers. We found that the cells located in the outer EGL were indeed mostly round and lacking long cellular processes (e.g. Fig. 4C-E). However, as shown in Fig. 3, PH3 positive cells located within the middle and inner EGL, where GCs undergo tangential migration, often possessed long and elaborate cellular processes, reminiscent of leading processes of migrating neurons (Fig. 3A). Additionally, in our time-lapse movies, we also observed cells extending long and motile processes that continuously changed their shape, length and direction (Fig. 3B). We therefore conclude that mitotically active GCPs can be located in the middle and inner EGL where tangentially migrating cells are found and can extend elaborate cellular processes before dividing.

Next, we characterised the morphologies of GCPs at different stages of mitosis, based on the pattern of PH3 staining. We found that the proportion of GCPs that extend processes decreases as the cells reach metaphase (Fig. 4A-E). Quantification in Fig. 4F shows that nearly all cells observed in prophase (96%) extended some form of a process, whereas very few cells in metaphase (23%) and no cells in anaphase or telophase extended processes. This suggests that dividing GCPs retract all their processes prior to anaphase. This conclusion was strengthened by our time-lapse movies where all observed dividing cells retracted their cellular processes and completely rounded up before division (Fig. 4G).

**Figure 3.**
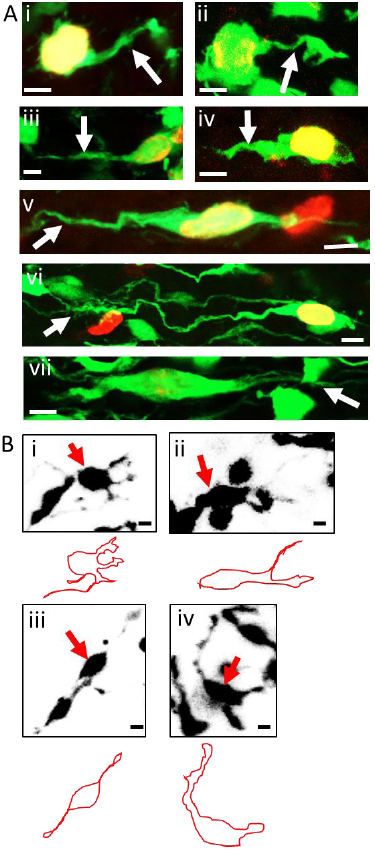
Granule cell precursors can extend elaborate processes before division. **A**) High magnification composite images of individual GFP (green) labelled cells that co-label with PH3 antibody (red) and possess a process. A variety of processes is observed (white arrows), some resembling leading processes (e.g. Av,vi). **B**) Stills of four different GCPs captured during time-lapse imaging of cerebellar organotypic slices. The cells (red arrows) extended different types of processes before mitosis. An outline of the cell is presented beneath the stills for better visualisation of the morphologies. Scale bar= 5 μm

Our time-lapse movies of the dividing GCPs allowed us to observe a number of other unexpected cellular behaviours. One representative example of a time-lapse imaging of a dividing cell in the EGL is shown in Fig. 5, where 12 stills are included, spanning 26 hours of imaging. The GCP imaged extends a leading‐ and a trailing-like processes before mitosis, its cell body migrates a short distance (∽10 μ m), followed by retraction of all processes, the cell rounding up and a parallel cell division taking place. Most divisions observed in the movies (64%) happened perpendicular to the pial surface with the rest being parallel (36%) (data not shown). Following division, the two daughter cells migrate in opposite directions from one another and extend their own processes, one forming a T-shaped cell process, reminiscent of forming parallel fibres. This type of behaviour was observed in the majority of imaged dividing GCPs, where, after division, the daughter cells extended leading processes in opposite directions and migrated away from each other (data not shown). Cells were not imaged long enough to follow the fate of the daughter cells.

GCPs undergo well characterised changes in gene expression as they differentiate in the EGL, with a number of markers associated with differentiating GCPs, including Axonin1 (TAG1 in mouse), which is important for axon pathfinding in the EGL^28^. We therefore stained the cerebellar tissue with anti-Axonin1 antibody, which was found in a broad domain in the lower half of the EGL (Fig. 6) and found that some PH3 positive cells co-localised with the Axonin1 expression domain in the inner EGL, suggesting that some GCPs can undergo mitosis even after their expression of postmitotic markers has commenced. TAG1 in the mouse^29^ and other postmitotic markers^30^ have previously been reported to co-label with proliferative markers in GCs in the middle EGL, suggesting an intermediate progenitor state.

**Figure 4.**
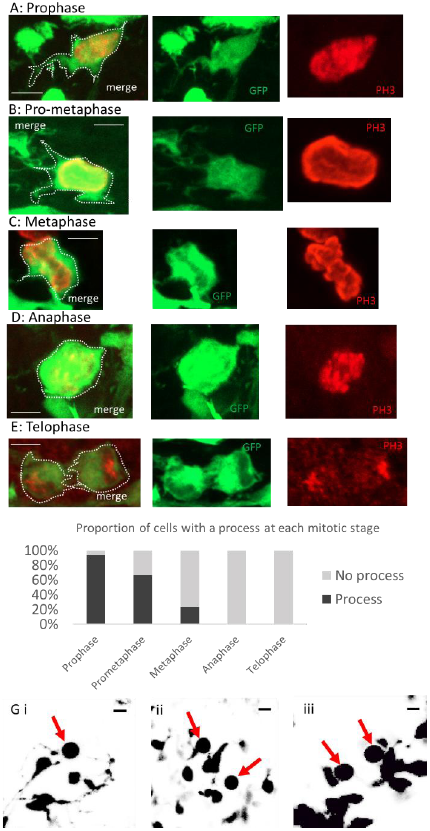
Granule cell precursors retract all processes before cell division. **A**) Examples of typical cellular morphologies of GCPs at different stages of mitosis. GFP signal (green), PH3 label (red) and a merged image are presented. White dots delineate the cell morphology. **A)** A cell in prophase with an extended process. **B)** A cell in pro-metaphase with processes that appear to be retracting. **C)** A cell in metaphase with very short processes. **D)** A cell in anaphase with no processes visible. **E)** A cell in late telophase. There are no processes extended at this point. **F)** Numbers of cells with extended processes at different stages of mitosis. As the mitotic cell reaches anaphase, it is increasingly less likely to be extending a process. **G)** Stills of five different GCPs captured during time-lapse imaging of cerebellar organotypic slices. The cells (red arrows) have no processes and a completely round morphology. The stills are shown just before the cells underwent division. All observed cells in time-lapse imaging underwent this stage of morphological transition before division. Scale bar=5 μm

NeuroD1 is a key differentiation marker of GCPs expressed in postmitotic neurons in the inner EGL^31–34^, even though its expression has been reported in mitotic GCs in some studies^35^. We therefore tested whether misexpression of NeuroD1 in GCPs will force them out of proliferation. To this end, we electroporated a plasmid encoding a full-length NeuroD1 conjugated to GFP at E6, the time of GCP birth at the rhombic lip, and observed the effect on proliferation two days later, at E8, in the forming EGL. We observed that many GCPs misexpressing NeuroD1 nevertheless express PH3 at E8 and therefore are able to undergo mitosis (Fig. 7A-B). However, misexpression of NeuroD1 leads to premature differentiation of GCPs and a depleted EGL (Fig. 7C-D) and results in a smaller and unfoliated cerebellum at E11 (Fig. 7E-G). Therefore, premature expression of a postmitotic marker in GCPs does not lead to immediate cessation of mitotic activity yet the presence of differentiation factors can decrease the proliferative potential of progenitor cells.

## Discussion

Cerebellar GCs are traditionally viewed to undergo well characterised sequential changes in morphology as they transition through the layers of the developing cerebellum. Here, we present evidence that the morphological changes of mitotic GCPs are complex and include more diverse cellular features than previously described. Using immunohistochemistry on fixed chicken cerebellar tissue, as well as imaging living cells undergoing mitosis in cerebellar organotypic slices, we demonstrate that GCPs divide in all layers of the EGL, can extend elaborate processes in between cell divisions, retract all processes prior to cytokinesis, and are able to undergo short migration movements before mitosis. Additionally, GCPs can divide perpendicular or parallel to the pial surface and the daughter cells tend to migrate away from one another in the medio-lateral direction. Therefore, the sequence of morphological differentiation of GCPs does not fully follow the linear sequence anticipated from Golgi studies. We propose that GCPs lose their proliferative potential as they migrate through the layers of the EGL, as depicted in the model in Fig. 8A. Mitotic granule cell precursors can express a number of ‘postmitotic’ markers such as Axonin1/TAG1^30^ and NeuroD1, as well as p27^29^. Interestingly, both their morphological transitions and the gradual transcriptional changes resemble those of the neuroblasts migrating within the RMS^19–22^. RMS neuroblasts go through repeated stages of migration, process retraction, and division on route to the olfactory bulb^19^. They also have a transcriptome that gradually shifts from a program controlling proliferation to one that modulates migration and differentiation^4^. While migrating in the RMS, neuroblasts receive a plethora of stimuli that modify transcription according to the local microenvironment. It would be interesting to see whether GCPs, which migrate tangentially within the EGL for shorter distances^36^ also respond to local stimuli to guide their development and what the identity of such stimuli is. The gradually shifting transcriptome changes are illustrated in Fig. 8B and include many other factors that each contributes to the cell fate decisions of GCPs. Exactly how these contributions are interpreted at the molecular level to result in a cell fate decision await further clarification.

**Figure 5.**
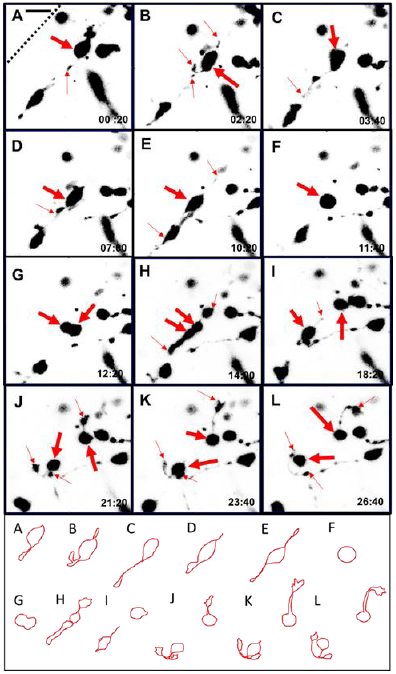
A time-lapse movie of a dividing granule cell precursor illustrates that they can migrate short distances before division after which their daughter cells migrate in the opposite directions. A selection of time-frames from a time-lapse movie of cerebellar slices. Small red arrows point to processes that extend from the cell body. Thick red arrows point to the cell of interest and its two daughter cells after division. The panel below shows a surface of each cell for visualisation. Dotted line in A represents the pial surface **A**) At the start of observation, the cell is extending one leading process. Whether it has a trailing process cannot be determined due to obstruction from another cell. **B**) Within 2 hours, the cell has migrated a small distance in a medio-lateral direction and extends two small, thin processes. The leading process seems to bifurcate and a growth cone is visible at the tip of one of the processes. **C**) The cell continues to extend its leading process, which has become longer and thinner. **D**) The cell has migrated further, and extends a thicker and shorter process. **E**) Within 3 hours the cell once again shows two long, thin processes from both sides prior to **F**) the cell retracting all processes and rounding up for cell division. **G**) within an hour of division, the cell divides perpendicularly to the pial surface. **H**) Both daughter cells extend their own processes in the opposite directions. One daughter cell extends a process with a very broad growth cone, whereas the other daughter cell has a longer and much thicker process. **I**) The daughter cells separate from each other as each migrates in a different direction. **J-L**) One of the daughter cells seems to extend two processes that bifurcate into a T-shaped process, resembling formation of a parallel fibre. The other daughter cell extends a long process towards the pial surface with a large, very motile growth cone. At the end of the time-lapse, the cell seems to turn its process back towards the inner EGL. Scale bar = 20μm

**Figure 6.**
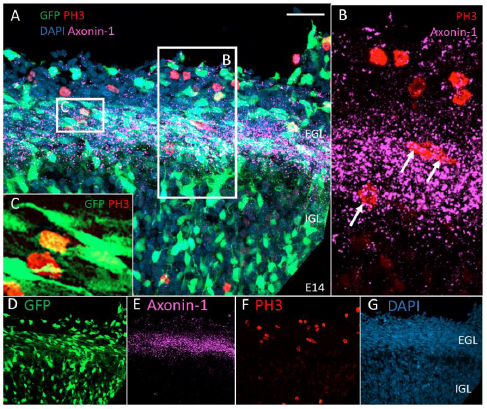
PH3 labelled cells in the EGL can co‐ label with a postmitotic marker Axonin1/TAG1. Cerebellar tissue from chick embryos electroporated with Tol2:GFP at E4 and fixed at E14 was stained for PH3 (red) and Axonin-1 (magenta). **A)** An example of cerebellar tissue showing GFP expressing cells, PH3 staining cells, Axonin-1 expression and DAPI stain. **B)** A magnified view of the area in A showing PH3 and Axonin-1 expression only. PH3^+ve^ cells are located in the Axonin-1^+ve^ area (arrows). **C)** A magnified view of the boxed area in A showing GFP expressing cells labelled with PH3. The cell is surrounded by differentiated cells with long processes and is located in Axonin-1^+ve^ area. **D-G)** Individual channels from A. **D** GFP, **E** Axonin-1, **F** PH3 and **G** DAPI staining. Scale bar= 50μm

Basal attachment has been proposed to be essential for GCPs proliferation^12,37,38^ however results obtained in this study clearly demonstrate that many GCPs divide without a basal attachment. Nevertheless, many cells with a basal attachment were observed in the oEGL in fixed cerebellar tissue (data not shown), resembling cells seen in the Cajal drawings. One possibility is that the attachment to the basal lamina is necessary for the expression of Math1, a transcription factor essential for transit amplification and symmetrical proliferative divisions of GCPs. As GCPs leave the outer EGL, they might lose their proliferative capacity and undergo asymmetrical or terminal neurogenic divisions only. Whether cell fate decisions correlate to the basal attachment of GCPs remains to be explored.

## Methods

### Chicken *in ovo* electroporation

Fertilised chicken eggs were incubated at 38 ^º^C for 4 days. The eggs were drained a day before use with a hypodermic needle to remove 3‐ 4ml of egg white. The eggs were then windowed using egg scissors and the embryo was located and its position manipulated for ease of access. DNA constructs at 1-2μg/μl were mixed with trace amounts of fast green (Sigma) and injected into the fourth ventricle directly below the rhombic lip using a glass needle. Where two or more constructs were co-electroporated both were mixed in equal concentrations, each at concentration of at 1-2μg/μ1. The negative electrode was placed underneath the embryo at the level of rhombic lip and the positive electrode was placed on the embryo at the same level to target the cerebellar rhombic lip. Three 50ms/10V square waveform electrical pulses were passed between the electrodes so that DNA entered the right side of the neural tube. Eggs were then treated with the Tyrode’s solution (1ml) and sealed back using tape. Eggs were incubated at 38°C until the embryos were harvested at appropriate experimental age (E8-E14). Whole embryos or their dissected cerebella were fixed overnight at 4^º^C in 4% PFA in PBS.

### Plasmids

pT2K-CAGGS-EGFP and pCAGGS-T2TP (together referred to as Tol2:GFP) were described previously^39^. pT2K-CAGGS-EGFP is a Tol2 transposon-flanked EGFP and pCAGGS-T2TP codes for Tol2 transposase. Upon the co-electroporation of pT2K‐ CAGGS-EGFP and pCAGGS-T2TP into a cell, the resulting transposon construct is excised from the plasmid and integrated into the host genome in proliferating precursors.

**Figure 7.**
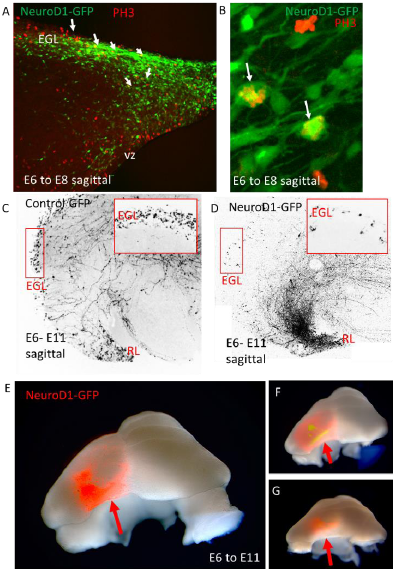
NeuroD1-misexpressing GCPs are able to undergo mitosis but have a limited proliferative potential and differentiate early. **A)** A plasmid encoding full length NeuroD1 and GFP was electroporated into E6 embryos which were fixed two days later, at E8, and stained for PH3. A sagittal cut through the forming EGL reveals co-expressing cells (arrows). B) Higher magnification of EGL cells co-expressing NeuroD1‐ GFP and PH3 (arrows). C and D) Embryos were electroporated with a control GFP plasmid (C) or NeuroD1‐ GFP (D) and fixed five days later, at E11. In the control electroporation there is a thick EGL (box, insert) whereas after NeuroD1 misexpression, the EGL is nearly devoid of cells (box, insert), which are located in the IGL close to the rhombic lip (RL). E-G) Three examples of whole cerebella electroporated with NeuroD1-GFP on one side of the rhombic lip only (red arrow). The electroporated side of the cerebellum shoes reduced size and lack of foliation pattern seen on the unelectroporated side. Scale bar= A,C,D= 100 μm B= 10 μm

NeuroD1 overexpression construct was cloned by T. Butts using full length NeuroD1 sequence following a β-actin promoter. IRES sequence was inserted between the *NeuroD1* sequence and the sequence for EGFP. A CAGGS-GFP plasmid was used as a control.

### *Ex vivo* time-lapse imaging

Chicken embryos were electroporated with Tol2:GFP constructs at E4 and the embryos were incubated up to embryonic day 11 to 14. Cerebella were then dissected in cold HBSS and their fluorescence level was determined using an epifluorescent microscope. Cerebellum with appropriate levels of fluorescent cells was chopped using a tissue chopper into 250mm slices. Slices with best cellular morphologies were chosen and transferred using a plastic pipette into a pre-assembled coverslip with a glass ring attached with silicon grease to create vacuum in the Rose chamber. The chamber is constructed from two 25 mm2 coverslips, a silicon spacer, a metal planchet milled to accept a condenser lens. The whole assembly is held together by two metal clips attached to the sides. The chamber can be filled and drained using two 25G needles and a syringe. Excess liquid transferred with the tissue was removed from the cover slip, making sure that the cerebellar slice lies flat on the cover slip. 500μl of rat tail collagen prepared to a neutral pH was then added on top of the slice in the glass ring, making sure that the slice remains close to the surface of the cover slip. The cover slip with the cerebellar slice was then incubated at 37^º^C 5% CO_2_ for 30 min to 1hr. After the collagen has set, another cover slip was placed on top of the ring and the rest of the Rose chamber was assembled. 2ml of pre‐warmed (37^º^C 5%CO_2_) culture medium (Basal Medium Eagle, 0.5% (w/v) D-(+)-glucose, 1% B27 supplement, 2 mM L-Glutamine, 100 U/mL penicillin, 100 μg/mL streptomycin) was added to the sealed chamber and the preparation was immediately imaged using a Nikon Eclipse EZ-C1 confocal microscope with a 20x objective lens overnight (12-28hrs) with 20min intervals between time points. For analysis, variable z-stack projections were chosen from the whole z‐ stack, depending on the best combination to observe specific cell morphologies.

**Figure 8.**
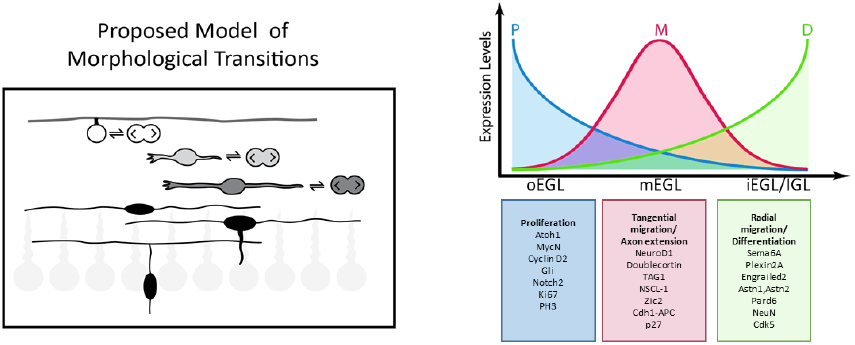
Proposed new model of GCP differentiation in the EGL. **A)** The current model of GCP morphological transitions postulates that mitotically active, polyhedral or round cells, with a possible basal attachment to the pial surface proliferate in the outer layer of the EGL. Only after the cell becomes postmitotic, it extends small horizontal processes and begins tangential migration. The processes continuously extend until the cell makes a switch to radial migration, at which point the horizontal processes are considered to be nascent parallel fibres. We suggest that this model is incomplete. The proposed model retains and confirms the morphological features of GCs in the EGL, but suggest that cells previously considered postmitotic, can in fact be proliferative. Divisions of GCPs occur in all layers of the EGL and cells with long horizontal processes are able to undergo mitosis by retracting all their processes and rounding up before division. In the proposed model, white shading represents a highly proliferative precursor, and black shading denotes a postmitotic cell. As cells migrate closer towards the inner EGL, the proliferative potential of the cells decreases, which means that the number of times the cell divides differs in the different layers, with many mitoses in the oEGL and few mitoses in the iEGL. (Purkinje cells are shown in light grey.) **B)** The suggested transcriptome of developing granule cells resembles the one found in SVZ‐derived neuroblasts in the RMS where their gene expression gradually shifts hence proliferation and differentiation genes can overlap. Genes responsible for granule cell precursors proliferation are most highly expressed in the outer EGL and are continuously downregulated as the cells migrate through the EGL layers and towards the IGL. Genes involved in tangential migration are upregulated in the middle and inner EGL. Differentiation-related genes start to be expressed in the middle and inner EGL and are strongly upregulated during radial migration and final stages of differentiation in the IGL.

### Histology and Immunohistochemistry

E14 chicken cerebellum was fixed in 4% PFA/PBS overnight. The tissue was then washed in PBS 3 times for 15 min and transferred into 10% sucrose (Sigma) in PBS for 30min until the tissue was perfused. The tissue was then transferred to 20% sucrose solution until perfused and finally was transferred into 30% sucrose solution and allowed to perfuse overnight. The tissue was then transferred into OCT compound (VWR) in moulds and placed on dry ice or liquid nitrogen to freeze. The tissue was stored at -80˚C overnight. For sectioning, the blocks were placed at -20˚C an hour before cutting to raise their temperature. The tissue was mounted on cryostat chucks using OCT compound and cut using a Zeiss Microm HM 560 cryostat at 50μm thickness and transferred onto Superfrost Plus slides (VWR). The cut sections were allowed to air dry for two hours and were stored at -80˚C long term and -20˚C short term. Cryostat sections were defrosted for at least 30min at room temperature. The slides were then washed three times for 5 min in PBS. Slides were then covered in 500-800μl blocking solution (1% normal goat serum, 0.2% Triton in PBS) and incubated for 30min at RT. Primary antibody was diluted at an appropriate concentration in the blocking solution. After the blocking solution was removed from the slides, 150-200μl of the antibody solution was added onto the slide and covered with parafilm to prevent drying. Incubation was performed overnight at 4^º^C. The next day, primary antibody was washed off with PBS three times for 5 mins. Secondary antibody was diluted in block solution and put onto the slides for 2hrs at RT. The slides were then washed with PBS three time for 5mins and covered with a coverslip using Fluoroshield mounting medium with DAPI (Abcam). Fluorescent confocal images were taken with Zeiss LSM 800 microscope. Z‐ stack projections were compiled using ImageJ. Z-stacks were taken at 1-202μm intervals.

